# Hyaluronic Acid Plays Differential Molecular Weight and Concentration Dependent Pathway Centric Changes to Human Lung Derived Microvascular Endothelial Cells in Culture

**DOI:** 10.64898/2026.06.01.729187

**Authors:** Shushruth Yellumahanthi, Kyoko Kojima, James A. Mobley

**Affiliations:** Cancer Cell Biology Masters Program, University of Alabama at Birmingham, Birmingham, AL, USA; Institutional Research Core Program, University of Alabama at Birmingham, Birmingham, AL, USA; Department of Anesthesiology and Perioperative Medicine, The Heersink School of Medicine, University of Alabama at Birmingham, Birmingham, AL, USA

**Keywords:** Hyaluronic Acid, Pulmonary, Human Microvascular Endothelial Cells (HMVEC), Mass Spectrometry, Proteomics, Systems Biology

## Abstract

**Background:** Hyaluronan (HA) is a major extracellular matrix glycosaminoglycan that regulates vascular integrity and immune signaling in the lung. Its biological effects are strongly size-dependent, with high-molecular-weight HA (HMW-HA) generally protective and low-molecular-weight HA (LMW-HA) pro-inflammatory. However, how different HA sizes and concentrations globally remodel endothelial cell signaling remains poorly understood.

**Methods:** Human lung microvascular endothelial cells (HULEC-5a) were treated with physiologic (200 ng/mL) or supraphysiologic (1 µg/mL) concentrations of LMW-, medium-molecular-weight (MMW-), or HMW-HA. Cell viability was confirmed by LDH assay. Quantitative proteomics with downstream Ingenuity Pathway Analysis (IPA) was used to profile HA-induced signaling networks.

**Results:** Proteomic analysis revealed a conserved HA-response signature across all conditions involving cell cycle regulation, senescence, and immune modulation, with distinct size-and dose-dependent differences. At supraphysiologic concentrations, HMW-HA suppressed proliferative and inflammatory pathways, consistent with a protective, quiescent phenotype. LMW-HA induced the broadest stress-associated proteomic changes, consistent with its role as a damage-associated molecular pattern.

Unexpectedly, physiologic MMW-HA elicited the strongest responses, driving metabolic and cytoskeletal pathways including insulin signaling and Rho GTPase activity. Network analysis highlighted 176 overlapping pathways across HA treatments, with unique contributions of LMW- and HMW-HA to stress- versus barrier-stabilizing signaling, respectively.

**Conclusion:** HA is not a passive structural molecule but an active regulator of endothelial signaling, with effects shaped by both molecular weight and concentration. Our findings identify a distinct role for MMW-HA at physiologic levels and highlight how HA fragmentation and accumulation may contribute to endothelial dysfunction in lung injury, with implications for targeted HA-based therapies.

## Introduction

Hyaluronan (HA), also historically known as hyaluronic acid, was first isolated in 1934 from the vitreous humor by Karl Meyer and John Palmer. Since then, it has been recognized as a ubiquitous glycosaminoglycan present in connective, epithelial, and neural tissues. Beyond its fundamental roles in structural integrity and cell signaling, HA has been widely studied for therapeutic applications, ranging from osteoarthritis treatment to cosmetic dermatology (9,11,15). This broad biological and clinical relevance underscores the importance of understanding HA’s diverse functions in health and disease (9,15).

Hyaluronan (HA) is a ubiquitous extracellular matrix glycosaminoglycan that plays critical structural and signaling roles in the lung. It is composed of repeating disaccharides (D-glucuronic acid and N-acetylglucosamine) and exists predominantly as a high-molecular-weight polymer in healthy tissues (1,2,9,15). In the lung, abundant HA is found in the peribronchial and interalveolar interstitium and as a key component of the endothelial glycocalyx (2,9). The endothelial glycocalyx is a meshwork of HA, proteoglycans, and glycoproteins on the luminal surface of blood vessels that preserves vascular homeostasis (2,9). By forming a hydrated, gel-like layer, HA contributes to the lung’s barrier function by regulating vascular permeability, shielding the endothelium from shear stress, and restricting leukocyte adhesion (2,9,12). Consistent with these properties, a healthy HA-rich glycocalyx supports pulmonary capillary integrity and limits edema formation. Conversely, injury or inflammation can lead to shedding of this HA-rich glycocalyx, with loss of endothelial barrier protection (2,19). Thus, HA is not only a passive structural element but also a dynamic regulator of vascular permeability and tissue integrity in the lung.

### HA Synthesis and Turnover by HAS and Hyaluronidases

The synthesis and degradation of HA are tightly controlled by specific enzyme families. HA is synthesized at the inner plasma membrane by three hyaluronan synthase isoforms – HAS1, HAS2, and HAS3 – which extrude the growing polymer into the extracellular space (1). Each HAS isoform has distinct enzymatic properties and produces HA chains of differing length. Notably, HAS1 and HAS2 predominantly generate high-molecular-weight HA (>500 kDa), whereas HAS3 synthesizes HA of lower molecular weight (often ≤500 kDa) (2,9,15). These differences can explain the varied sizes of HA observed in tissues and may underlie context-specific functions of HA matrices (2,15). HA production is upregulated by pro-inflammatory stimuli (e.g. TNFα, IL-1β, LPS) (2,9), and changes in HAS expression are linked to lung pathology. For example, in ventilator-induced lung injury, mice lacking HAS3 failed to accumulate LMW-HA and had reduced neutrophil infiltration compared to wild-type, implicating HAS3-driven HA fragments in the inflammatory response (2,14).

HA catabolism is mediated primarily by hyaluronidases, among which hyaluronidase-1 (Hyal-1) and -2 (Hyal-2) are most active in somatic tissues (2,9,15). Hyal-2 is a glycosylphosphatidylinositol-anchored enzyme on the cell surface that initiates HA cleavage in the extracellular pericellular space (9,15,18). In concert with its co-receptor CD44, Hyal-2 attacks HMW-HA and generates intermediate fragments (∼20 kDa) (2,9). These fragments are then internalized and delivered to endo-lysosomal compartments where Hyal-1 – an acid-active lysosomal hyaluronidase – further degrades them into oligosaccharides (2,9,15). Through the combined action of Hyal-2, Hyal-1, and possibly other hyaladherins, HMW-HA can be rapidly broken down into bioactive fragments. Six hyaluronidase genes exist (HYAL1-4, PH20, and a pseudogene), but Hyal-1 and -2 are considered the chief regulators of HA turnover in the lung (2,9). In addition to enzymatic catabolism, physical and chemical insults can depolymerize HA – reactive oxygen species and mechanical forces (e.g. ventilator-associated stretch) can fragment HA independent of enzymes (2,3,15). The balance between HAS synthesis and hyaluronidase (and ROS-mediated) degradation determines the steady-state molecular weight distribution of HA (1,2,15). Perturbations in this balance during disease often result in accumulation of aberrantly sized HA polymers.

### Molecular Weight-Dependent Functions of HA

It is well established that the biological effects of HA critically depend on its molecular weight. Native high-molecular-weight HA (HMW-HA, often >10^6 Da) is generally immunosuppressive or homeostatic, whereas fragmented low-molecular-weight HA (LMW-HA, <∼200–500 kDa) can act as a potent pro-inflammatory signal (3,9,15). HMW-HA predominates in healthy lungs and contributes to immune quiescence and barrier stability. For example, exogenous HMW-HA administration has been shown to inhibit inflammation in models of lung injury (3,11,12), and HMW-HA engages mechanisms that preserve tissue integrity (11,12). One major HMW-HA receptor is CD44, a broadly expressed cell-surface glycoprotein that anchors HA on cell membranes and transduces extracellular matrix cues (3,9). Cross-linking of CD44 by HMW-HA can activate anti-inflammatory pathways – such as promoting the production of anti-inflammatory cytokines and scavenging of reactive oxygen species – and supports regulatory immune cells (3,9,11,12). HMW-HA-CD44 interactions also appear to attenuate Toll-like receptor (TLR) signaling. Indeed, HMW-HA can antagonize TLR4-mediated inflammatory responses; in mice, pretreatment with HMW-HA blunted LPS-induced TNFα/IL-6 production in a TLR4-dependent fashion (3,11,12). These observations have led to the concept that intact HMW-HA functions as a “tissue integrity” signal that actively dampens inflammation when the matrix is unperturbed (3,9,11,12).

By contrast, when tissue damage or stress causes HA to be depolymerized, the resulting LMW-HA fragments serve as endogenous danger signals. LMW-HA is recognized as a damage-associated molecular pattern (DAMP) that can trigger innate immune receptors analogous to pathogen-associated molecules (3,9,13). Fragments of HA engage pattern recognition receptors like TLR2 and TLR4 on immune cells, often in cooperation with CD44, to initiate pro-inflammatory signaling cascades (3,9,13,21). LMW-HA stimulation leads to activation of NF-κB and other transcription factors, driving the expression of chemokines and cytokines such as IL-8, IL-6, TNFα, and IL-1β (3,13,21). These signals promote leukocyte recruitment and maturation (for instance, dendritic cell activation and neutrophil chemotaxis) and can induce endothelial activation and proliferation (3,13). Thus, fragmented HA effectively alerts the immune system to tissue injury and contributes to the inflammatory milieu (3,9,13,21). Notably, LMW-HA’s ability to induce inflammation has been documented in sterile injury settings (absence of infection), underscoring its role as an intrinsic mediator of tissue damage response (3,13,17). Several HA-binding receptors besides CD44 and TLR4 can modulate these effects – for example, the receptor for HA-mediated motility (RHAMM) and TLR2 have also been implicated in LMW-HA signaling (3,4,9,13). Overall, HMW-HA versus LMW-HA often exert opposing effects: HMW-HA is typically anti-inflammatory and barrier-protective, whereas LMW-HA is pro-inflammatory and can destabilize cellular barriers. In the context of endothelium, HMW-HA helps maintain vascular integrity, while LMW-HA tends to disrupt endothelial junctions and glycocalyx, increasing permeability (4,13).

### HA Fragmentation in Lung Injury and Inflammation

The size-dependent functions of HA are especially relevant in acute lung injury and inflammatory lung diseases. Under injurious conditions such as acute respiratory distress syndrome (ARDS), ventilator-induced lung injury, or oxidant inhalation, there is excessive HA turnover and shedding. ARDS is accompanied by a marked increase in total lung HA content, which has been correlated with worse gas exchange (oxygenation impairment) in patients (5,17). This increase reflects not just more HA production but also degradation of HMW polymers into smaller fragments. In the setting of severe infection (e.g. COVID-19 pneumonia) or trauma, abundant HA fragments have been detected in the airspaces and circulation (1,6,19). These fragments are thought to contribute to the “cytokine storm” and vascular leak characteristic of ARDS (5,6,19). In a recent example, COVID-19 patients were found to have high levels of circulating HA fragments that directly induce endothelial barrier dysfunction in vitro in a CD44- and Rho kinase-dependent manner (19). This finding highlights how HA breakdown products can impair pulmonary microvascular integrity, leading to edema and exacerbating lung injury.

Non-infectious insults show a similar pattern. During hyperoxic or chemical inhalation injury, HA in the lung extracellular lining fluid undergoes oxidative depolymerization. For instance, chlorine gas exposure causes HA degradation into <300 kDa fragments that initiate a cascade of inflammation and tissue hyper-responsiveness (7). In a mouse model of chlorine inhalation, the rise in LMW-HA in bronchoalveolar lavage fluid was associated with increased lung inflammation and airway hyperreactivity, whereas treatment with exogenous HMW-HA after exposure could ameliorate these effects (7,11). This supports the concept that restoring high-molecular HA or preventing its breakdown has therapeutic potential in acute lung injury by preserving the endothelial–epithelial barrier. Likewise, mechanical ventilation at high tidal volumes has been shown to elevate LMW-HA levels in the lung and promote neutrophil infiltration, a process blunted in HAS3-knockout mice that cannot efficiently generate HA fragments (4,14).

Viral lung infections also intersect with HA metabolism. Respiratory syncytial virus (RSV), a common cause of pediatric bronchiolitis and lung injury, induces remodeling of the pericellular matrix. During RSV infection, lung structural cells exhibit altered HA turnover – studies have observed decreased hyaluronidase expression leading to HA accumulation in the airways, accompanied by heightened leukocyte adhesion and lung inflammation (8). In mouse models, RSV triggers excess HA deposition in bronchial tissue and increased inflammatory cell recruitment (8,20). These data suggest that an inability to degrade HA (and consequent buildup of HA, potentially of intermediate size) can contribute to persistent inflammation in viral injury. On the other hand, other studies have noted that HA fragmentation can also occur in viral ARDS; for example, influenza and SARS-CoV-2 infections are associated with HA-rich, viscous edema in alveoli, implying ongoing HA degradation in the diseased lung (1,19). In sum, whether by overproduction of HA or by excessive fragmentation (or both), a dysregulated HA matrix is a hallmark of lung injury. The fragmented HA acts as a pro-inflammatory mediator and disrupts endothelial and epithelial barrier function, whereas intact HMW-HA helps maintain vascular and alveolar integrity. This dichotomy underlies the need to understand how different sizes of HA influence lung cells, particularly the endothelium which is a key target in vascular leak and inflammation.

### Endothelial Cell Model and Proteomic Approach

Pulmonary microvascular endothelial cells are central to the pathology of lung injury, as they constitute the barrier that prevents vascular leakage into airspaces. To investigate how HA of different molecular weights modulates endothelial biology, we utilized the HULEC-5a cell line as an in vitro model of human pulmonary microvascular endothelium. HULEC-5a cells are immortalized (SV40-transfected) human lung microvascular endothelial cells that retain many characteristics of primary pulmonary endothelium (10,16). They form cobblestone monolayers and have been widely used to study lung endothelial function and dysfunction in diseases such as ARDS, infection, and pulmonary inflammatory disorders (10,16). This model allows us to probe endothelial responses under controlled conditions and is cost-effective and reproducible compared to primary cell isolations.

In this study, we exposed HULEC-5a monolayers to HA of three distinct size ranges – low molecular weight (LMW), medium molecular weight (MMW), and high molecular weight (HMW) – at two physiologically relevant concentrations. The “physiologic” concentration (∼200 ng/mL) approximates baseline interstitial levels of HA found in tissues or circulation, whereas a higher “supraphysiologic” dose (1 µg/mL) was chosen to mimic conditions of pathological HA accumulation (such as during severe inflammation or tissue injury) (5,19). After 24 hours of treatment, we assessed global protein expression changes in the endothelial cells using an unbiased proteomic approach. By quantitatively profiling the proteome, we aimed to capture the breadth of signaling pathways and cellular processes influenced by HA of varying sizes and concentrations. This approach is advantageous because HA can simultaneously impact multiple signaling axes – from inflammatory pathways (e.g. NF-κB, interferon signaling) to cytoskeletal dynamics (e.g. RhoA/ROCK pathway involved in barrier regulation) – and these may not be evident through targeted assays (6,19). Proteomic analysis enables the identification of both common responses (core pathways activated by HA in general) and divergent responses (unique effects of HMW-HA vs. LMW-HA, or high vs. low dose) in the endothelial cells. We hypothesized that HMW-HA, MMW-HA, and LMW-HA would differentially modulate endothelial signaling networks, with HMW-HA likely reinforcing barrier-protective or quiescent phenotypes and LMW-HA inducing pro-inflammatory or stress-related proteins, while intermediate-size HA might have overlapping or intermediate effects. The results of this investigation will provide a mechanistic framework for understanding how HA fragmentation state and abundance influence pulmonary microvascular endothelium. Such insights are important for high-impact pulmonary and vascular biology, as they could reveal novel targets for preserving endothelial function in lung injury (for instance, by supplementing HMW-HA or inhibiting HA fragment signaling) and improve our overall understanding of the endothelial glycocalyx and matrix signaling in lung health and disease.

Despite extensive work on HA biology, relatively few studies have systematically examined how HA of different molecular weights and concentrations shape endothelial signaling networks. A recent PubMed analysis highlights this gap: while tens of thousands of articles mention HA broadly, fewer than 15 directly address HA in the context of proteomics and the lung. This scarcity reflects a critical need for unbiased approaches to map how HA fragments versus intact polymers influence endothelial cells at a systems level. Addressing this gap will help clarify the dichotomy between “protective” high-molecular-weight HA and “damaging” low-molecular-weight HA, particularly under physiologic and supraphysiologic conditions relevant to lung injury.

## Methods

### Proteomics Analysis

Sample preparation for proteomics analysis was carried out on 20ug of protein from total cell lysate (MPER, ThermoScientific). GeLC analysis was performed with 5-MW fractions covering the entire lane for each sample. Mass spectrometry of the peptide digests was analyzed using an Agilent 1260 Infinity nHPLC system coupled in-line to a Thermo Orbitrap Velos Pro mass spectrometer operating in data-dependent acquisition (DDA) CID mode. Samples were separated on a Phenomenex C18 column using a multi-step binary gradient program and, following parent ion scans @ 60k resolution spanning 300-1200 m/z, fragmentation data were collected for the top 15 most intense ions. MS data conversion and searches were performed using XCalibur RAW data files, collected in profile mode, which was centroided and converted to MzXML format using ReAdW v3.5.1, and subsequently searched against a species-specific UniProtKB database using SEQUEST. For peptide filtering, grouping, and quantification, the list of peptide IDs generated from the SEQUEST search results were filtered using Scaffold (Protein Sciences, Portland, Oregon). Protein-level filters were set to >99.0% C.I. and a false discovery rate (FDR) <1.0. Relative quantification across experiments was then performed via spectral counting, and spectral count abundances then asses for potential outliers and normalized between samples prior to statistical analysis.

### Statistical Analysis

One-way ANOVA was utilized to compare multiple groups using Qlucore Omics Explorer to generate PCA plots and Heat Maps, in addition students t test was applied to the Volcano Plots. GraphPad Prism 8 was used to perform statistical analysis and create the graphs. Significance was defined as a p-value of less than 0.05. For the proteomic data, two separate statistical analyses were applied with a fold change cut-off of >|1.5|; nonparametric and parametric were performed for each pairwise comparison. The non-parametric analysis included: (1) calculation of weight values using Significance Analysis of Microarrays (SAM), with a cutoff of >|0.8|, and (2) a one-tailed, unequal variance t-test with a p-value cutoff of <0.05. The results were then sorted based on the highest statistical relevance within each comparison. Gene ontology assignments and pathway analysis were carried out using Ingenuity Pathway Analysis (QIAGEN, Redwood City, CA). Interactions identified within IPA were manually correlated using full-text peer-reviewed articles.

## Results

We first evaluated whether treatment with hyaluronan altered the viability of human lung microvascular endothelial cells (HULEC-5a). Lactate dehydrogenase (LDH) activity, a marker of membrane damage and cytotoxicity, was measured after 24-hour exposure to low-, medium-, or high-molecular-weight HA at physiologic (200 ng/mL) and supraphysiologic (1 µg/mL) concentrations. Across all conditions, LDH release remained comparable to vehicle-treated controls, indicating no evidence of cytotoxicity (Fig. 1). These results confirm that subsequent proteomic changes reflect specific biological responses to HA rather than cell injury.

**Figure 1.**
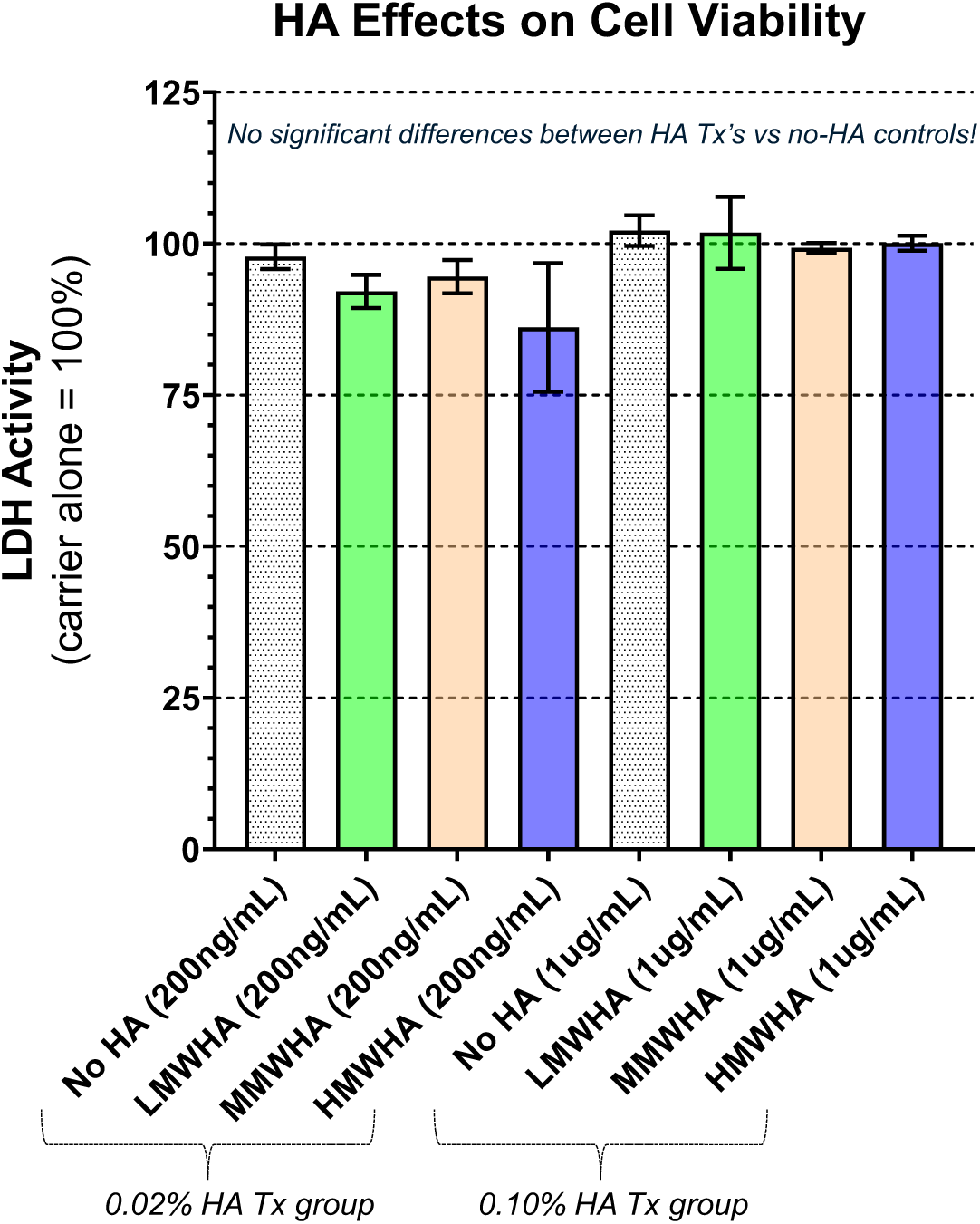
Effect of HA treatment on endothelial cell viability. HULEC-5a cells were treated with vehicle, LMW-HA, MMW-HA, or HMW-HA at physiologic (200 ng/mL) or supraphysiologic (1 µg/mL) concentrations for 24 hours. LDH activity was measured and normalized to vehicle control (set at 100%). No significant differences were observed across treatment groups (n=4 per condition).

We next investigated the effects of high-molecular-weight HA (HMW-HA) at a supraphysiologic concentration (1 µg/mL) on the endothelial proteome. Global proteomic profiling identified significant differences in protein abundance compared to untreated controls, with 158 proteins meeting criteria of *p* < 0.05 and fold change >1.5 (Fig. 2A, volcano plot). Principal component analysis and hierarchical clustering demonstrated distinct separation between HMW-HA–treated and control samples, indicating robust and reproducible treatment-dependent effects (Fig. 2B–C).

**Figure 2.**
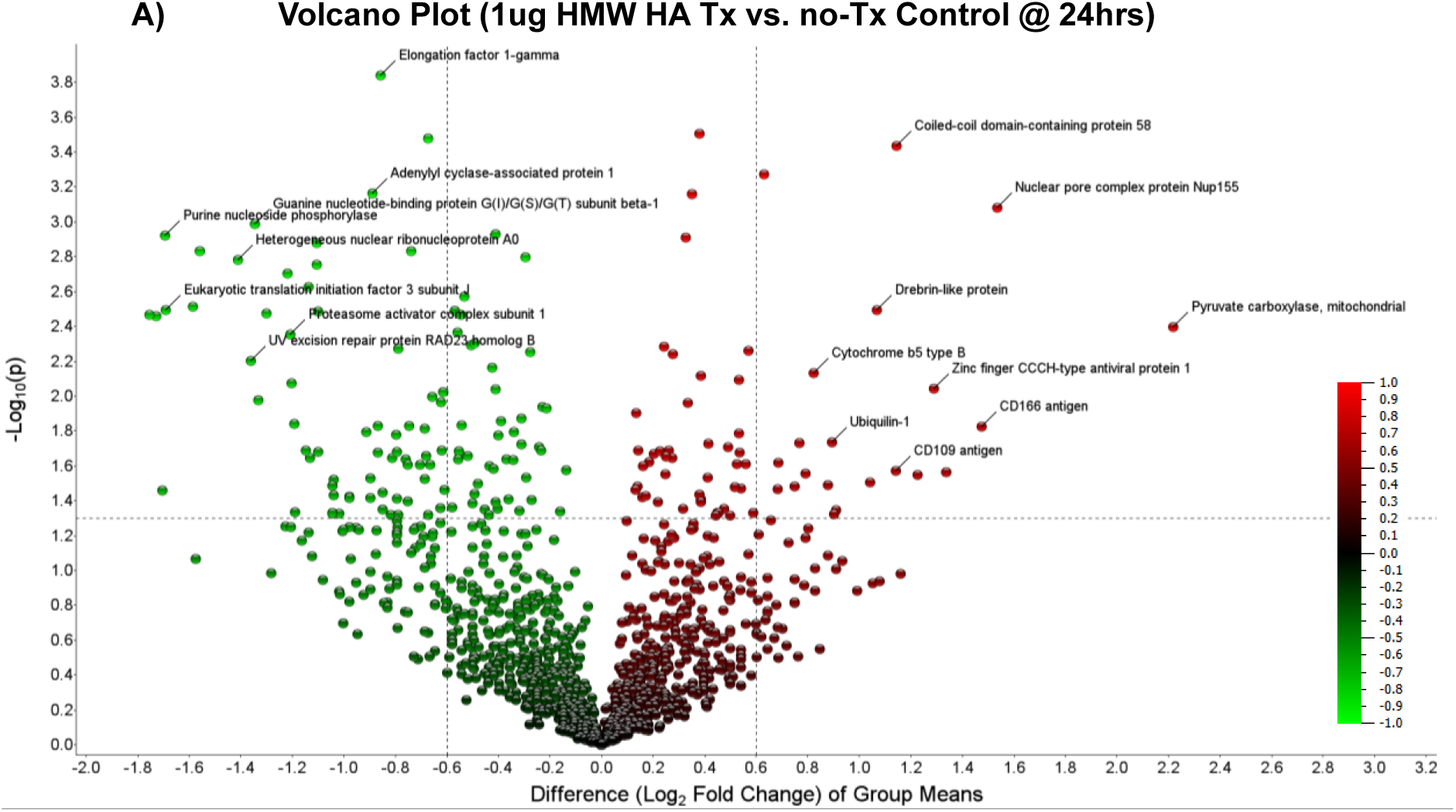
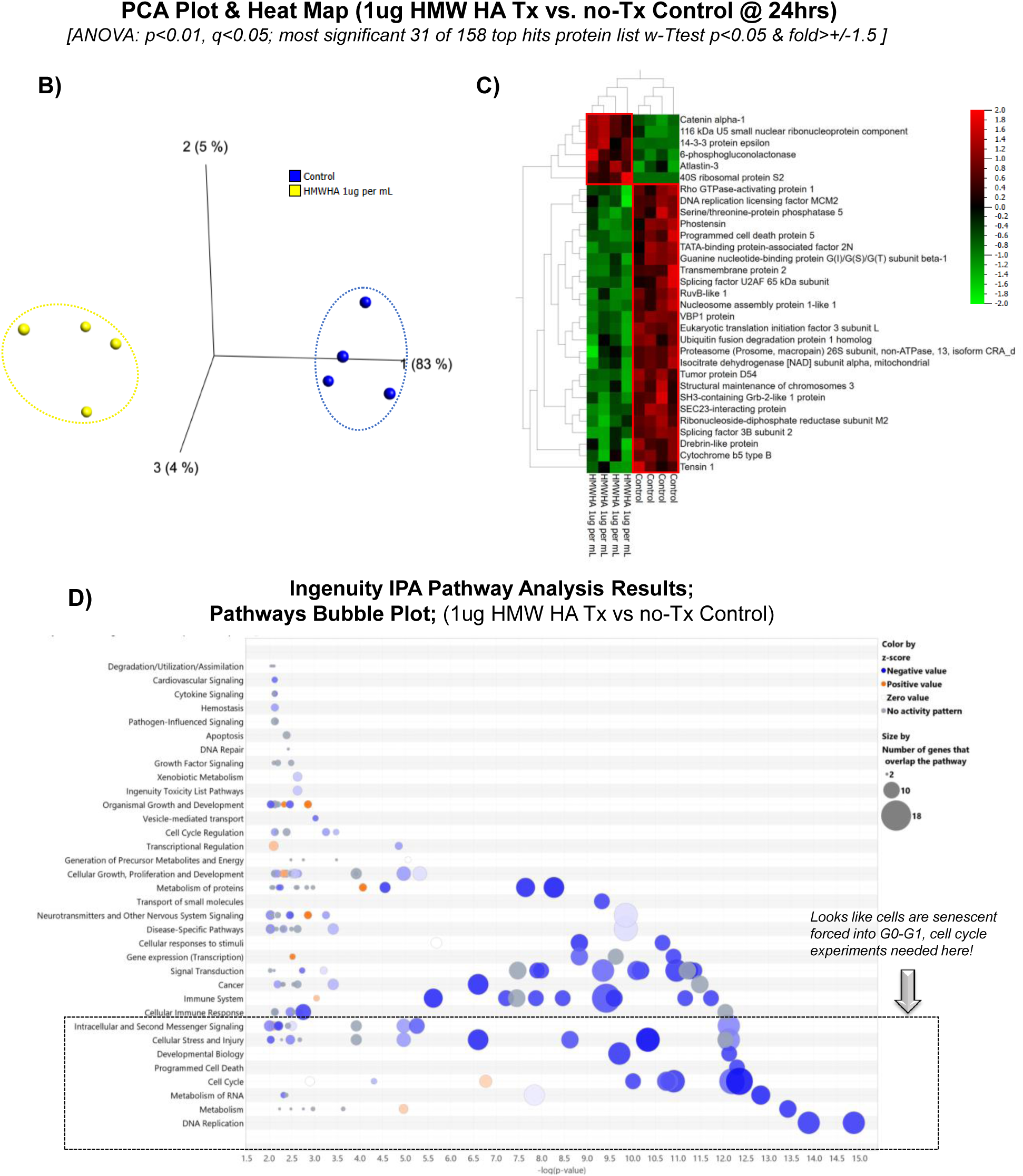
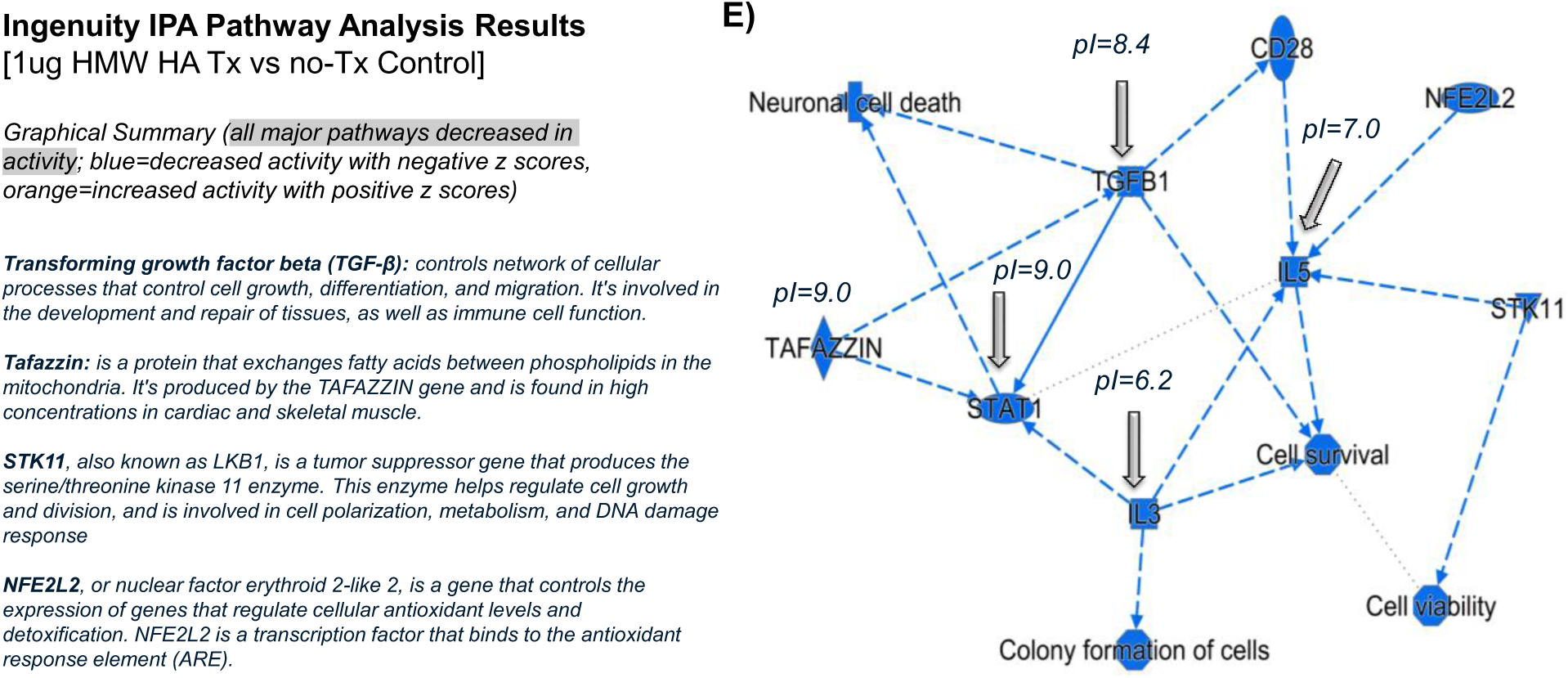
Proteomic effects of high-molecular-weight hyaluronan (HMW-HA) at supraphysiologic dose. HULEC-5a endothelial cells were treated with 1 µg/mL HMW-HA for 24 hours and analyzed by global proteomics. (A) Volcano plot of differentially expressed proteins versus untreated controls (p < 0.05, fold change >1.5). (B) Principal component analysis (PCA) and (C) hierarchical clustering heatmap demonstrate clear segregation of treatment groups. (D) Ingenuity Pathway Analysis (IPA) bubble plot and (E) graphical summary highlight widespread suppression of cell cycle and inflammatory pathways, consistent with induction of a protective senescent/quiescent phenotype.

Ingenuity Pathway Analysis (IPA) revealed widespread suppression of pathways related to cell cycle progression, DNA replication, and inflammatory signaling, with the majority of significantly altered pathways exhibiting negative z-scores (Fig. 2D–E). These changes are consistent with induction of an endothelial “protective senescence” phenotype, characterized by quiescence and reduced proliferative activity. Importantly, these proteomic shifts occurred in the absence of cytotoxicity, confirming that they represent active biological regulation rather than nonspecific injury.

To assess the influence of HA molecular weight on endothelial responses at supraphysiologic dose, we compared low-, medium-, and high-molecular-weight HA treatments (1 µg/mL each) against controls. Principal component analysis and hierarchical clustering revealed distinct proteomic signatures for each HA size, with clear segregation from controls and from one another (Fig. 3A–B).

**Figure 3.**
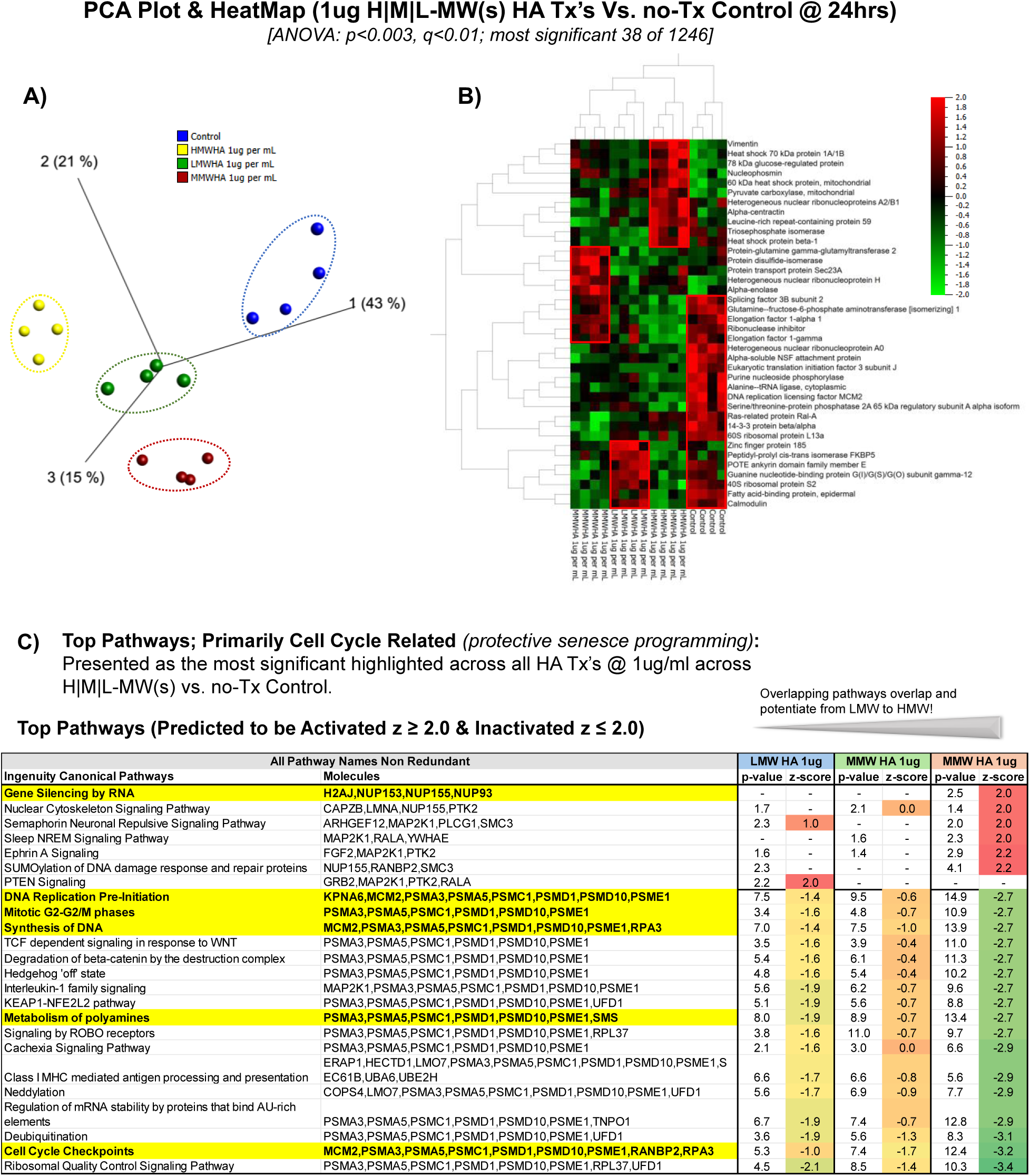
Comparative proteomic effects of low-, medium-, and high-molecular-weight hyaluronan (LMW, MMW, HMW) at supraphysiologic dose. HULEC-5a endothelial cells were treated with 1 µg/mL HA for 24 hours. (A) Principal component analysis (PCA) and *(B)* hierarchical clustering heatmap demonstrate distinct separation of HA-treated groups from controls and from each other. (C) Ingenuity Pathway Analysis (IPA) revealed extensive overlap across HA treatments (176 pathways), but with differences in magnitude: LMW-HA altered the greatest number of pathways, whereas HMW-HA produced stronger predicted inhibition of cell cycle and inflammatory signaling (z-score ≤ –2). MMW-HA exhibited intermediate activity.

Ingenuity Pathway Analysis identified a shared set of 176 pathways altered across all HA types, with consistent directionality of predicted activation or inhibition (Fig. 3C). Despite this overlap, notable differences in magnitude and scope were observed. LMW-HA altered the greatest number of pathways (301 total), followed by HMW-HA (272) and MMW-HA (intermediate). However, HMW-HA exhibited stronger predicted regulatory effects, with 60 pathways showing robust inhibition (z-score ≤ –2) compared to only 19 pathways for LMW-HA.

These findings suggest that while all HA molecular weights elicit a conserved endothelial response signature at high concentration, HMW-HA produces a more focused, potent suppression of cell cycle and inflammatory pathways, whereas LMW-HA drives broader but weaker modulation. MMW-HA displayed intermediate effects, indicating molecular weight governs both the breadth and strength of HA-induced proteomic remodeling.

We next examined the effects of HA at physiologic concentrations (200 ng/mL). Compared with untreated controls, all HA molecular weights produced detectable proteomic shifts, though the magnitude was reduced relative to supraphysiologic dosing. Principal component analysis and hierarchical clustering confirmed separation of HA-treated groups from controls, but with tighter clustering and smaller variance (Fig. 4A–B).

**Figure 4.**
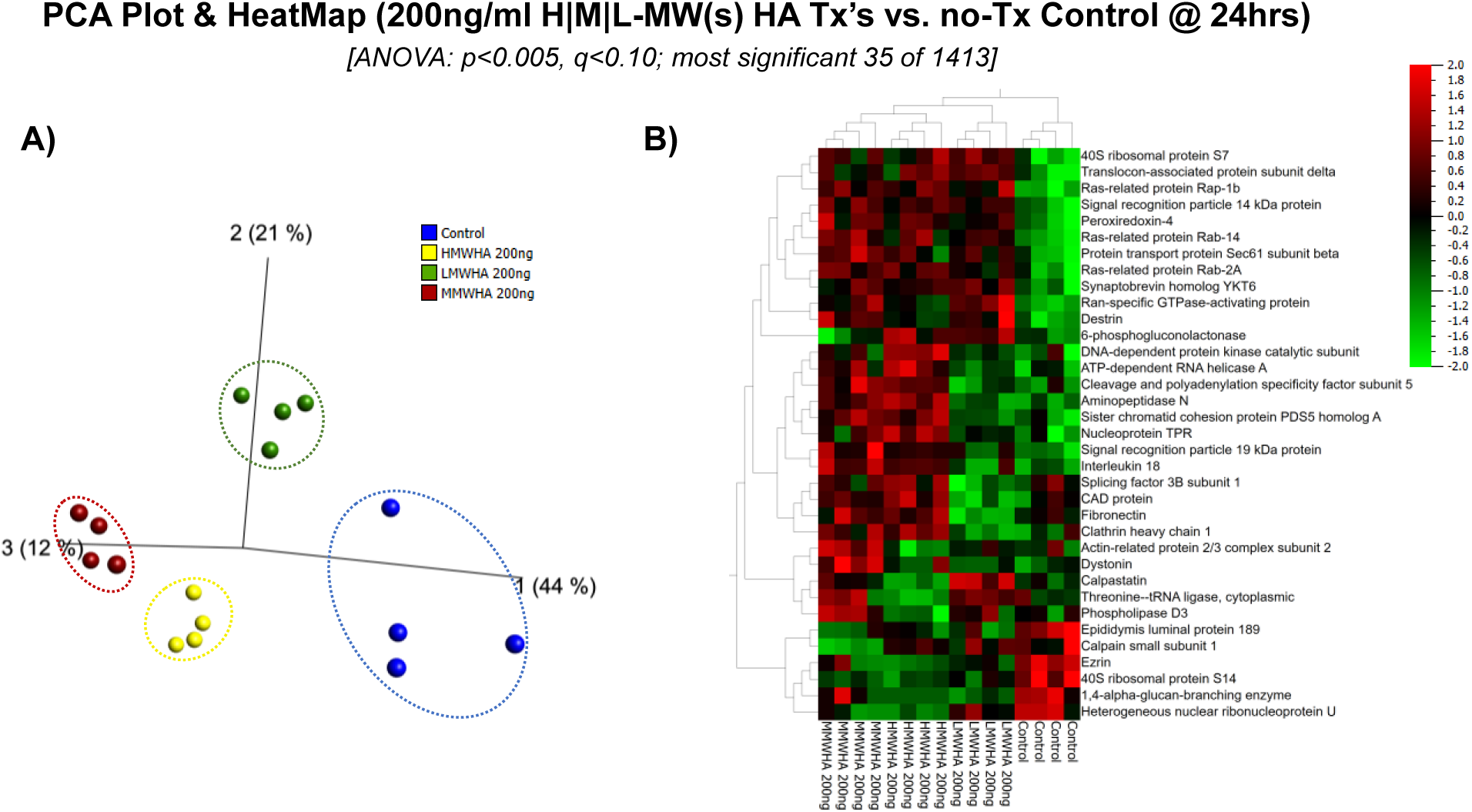
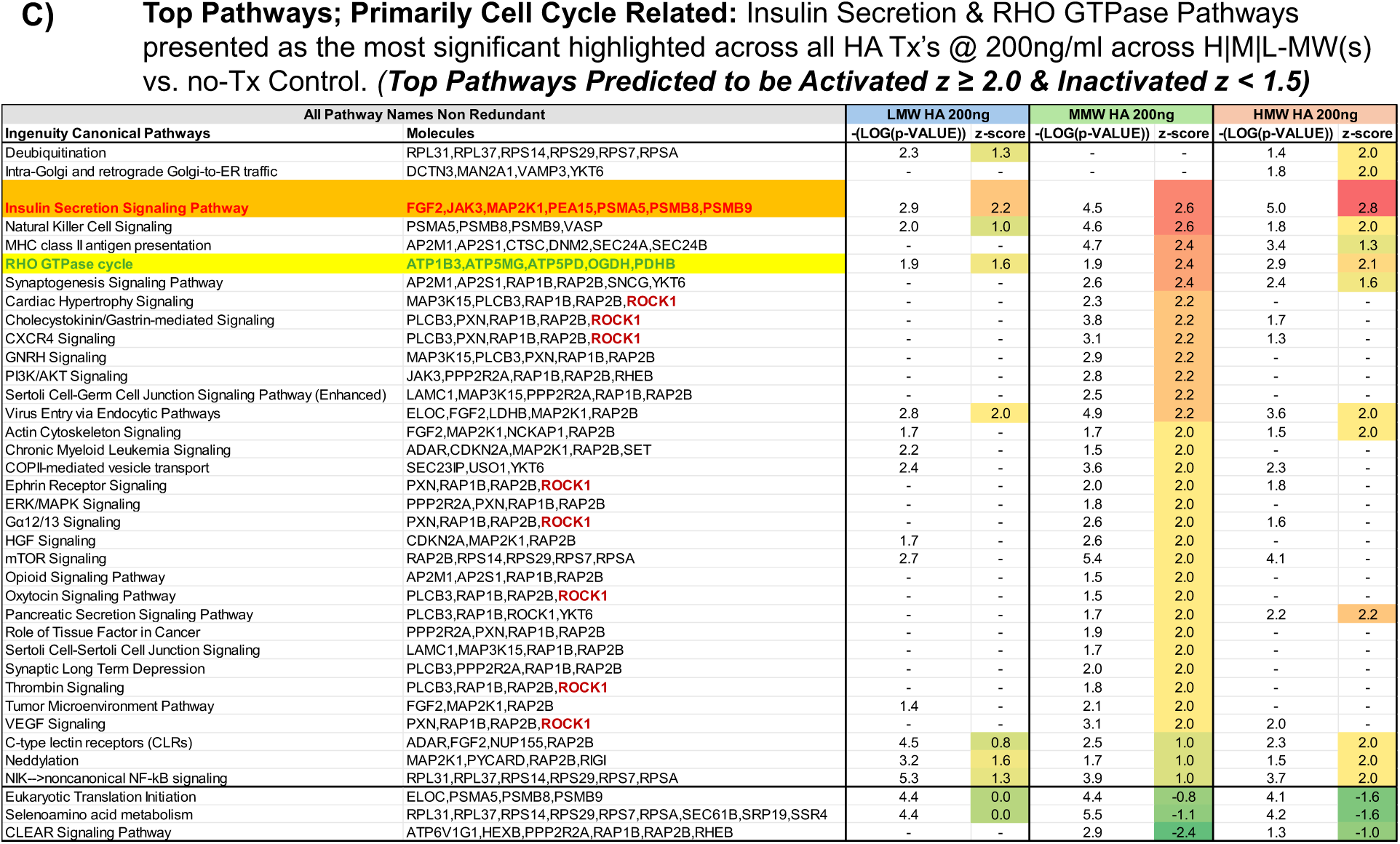
Proteomic effects of hyaluronan at physiologic dose. HULEC-5a endothelial cells were treated with 200 ng/mL of LMW-, MMW-, or HMW-HA for 24 hours. (A) Principal component analysis (PCA) and (B) hierarchical clustering heatmap demonstrate separation between HA-treated and control groups, though with smaller variance than at high dose. (C) Ingenuity Pathway Analysis (IPA) shows modulation of cell cycle, senescence, and immune pathways, with MMW-HA uniquely producing stronger predicted activation of insulin secretion and Rho GTPase signaling. These data indicate that MMW-HA is the most active HA form at physiologic concentrations.

Ingenuity Pathway Analysis revealed that many of the same pathways affected at high dose—cell cycle checkpoints, senescence, and immune signaling—were also modulated at physiologic levels, but with attenuated z-scores (Fig. 4C). Unexpectedly, medium-molecular-weight HA (MMW-HA) was significantly more active than either LMW- or HMW-HA, driving stronger predicted activation of pathways involved in insulin secretion and Rho GTPase signaling. These findings suggest that while all HA sizes influence endothelial proteomes at baseline levels, MMW-HA exerts disproportionate potency, particularly in metabolic and cytoskeletal pathways.

To obtain a global view of HA-induced changes, we performed network-level analyses across all molecular weights and doses. Network enrichment revealed a conserved set of pathways shared across treatments, including cell cycle regulation, deubiquitination/neddylation, immune suppression, and senescence signaling (Fig. 5A–B). These overlapping signatures suggest a core HA-responsive program in pulmonary endothelium.

**Figure 5.**
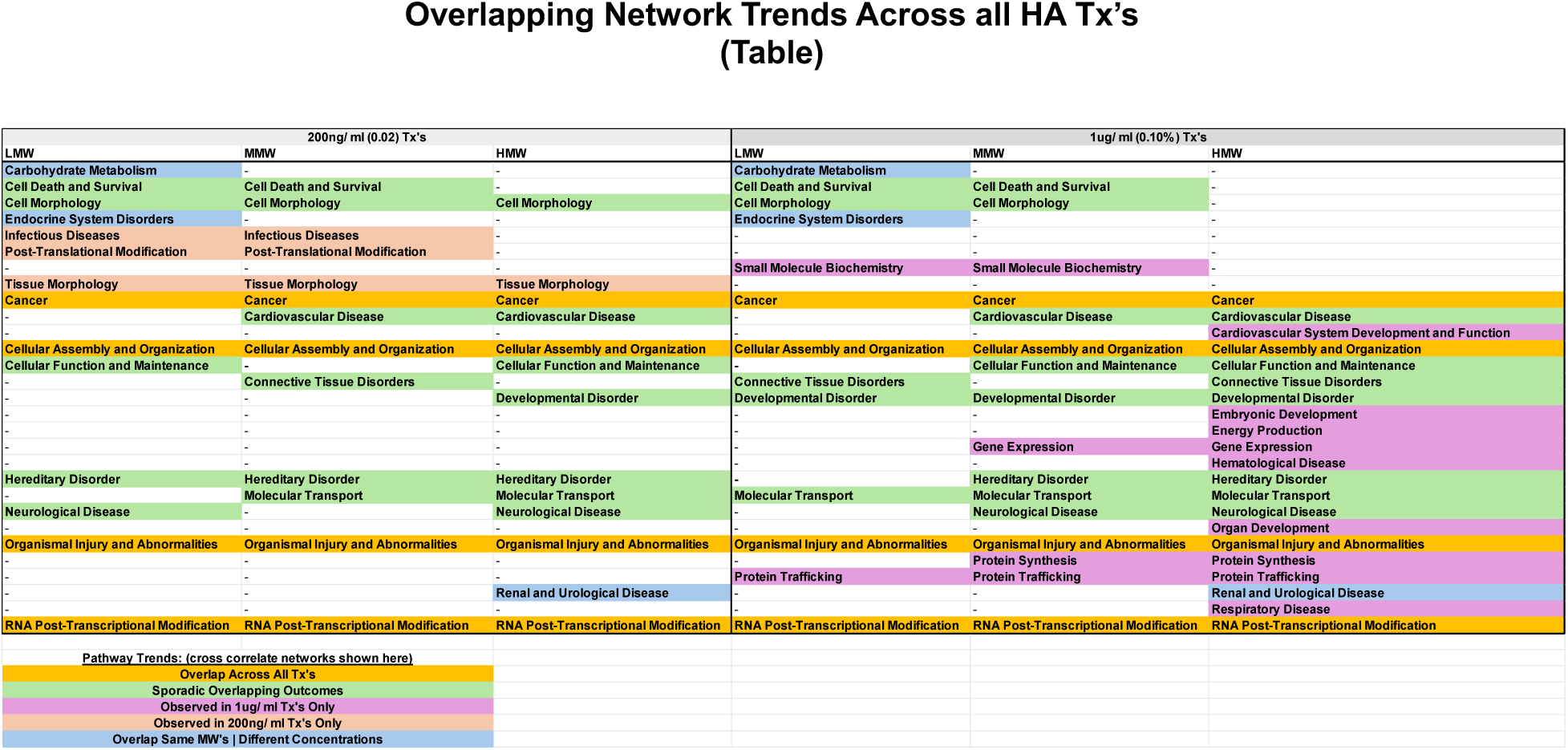
Overlapping network trends across hyaluronan (HA) treatments. Ingenuity Pathway Analysis (IPA) identified conserved and distinct biological networks altered in endothelial cells exposed to low-molecular-weight (LMW), medium-molecular-weight (MMW), and high-molecular-weight (HMW) HA at physiologic (200 ng/mL) and supraphysiologic (1 µg/mL) concentrations. The table highlights network categories that were shared across all HA treatments (e.g., cell morphology, cancer biology, cardiovascular disease), as well as those unique to specific doses or molecular weights (e.g., infectious disease networks at physiologic doses, respiratory disease and protein synthesis/trafficking at supraphysiologic doses, carbohydrate metabolism with LMW-HA, renal/urologic disease with HMW-HA). These patterns indicate that HA exerts both common and size- or dose-specific influences on endothelial signaling.

At the supraphysiologic dose (1 µg/mL), network activity was predominantly suppressed, with HMW-HA exerting the most potent downregulation of proliferative and inflammatory networks. In contrast, physiologic dosing (200 ng/mL) elicited weaker responses that trended toward homeostatic buffering, with MMW-HA producing uniquely strong modulation of insulin secretion and Rho GTPase pathways.

While LMW-HA induced the largest number of network alterations overall, its effects were diffuse and enriched in stress-related and metabolic categories, consistent with a damage-associated profile. By comparison, HMW-HA exerted more selective, endothelial-stabilizing effects, and MMW-HA emerged as the dominant regulator at physiologic dose. Together, these analyses highlight both the conserved and context-specific features of HA signaling, underscoring the importance of molecular weight and concentration as key determinants of endothelial responses.

## Discussion

We demonstrate here that hyaluronan (HA) of different molecular weights and concentrations induces distinct proteomic programs in pulmonary microvascular endothelial cells. Using unbiased quantitative proteomics, we identified a conserved HA-response signature encompassing cell cycle regulation, senescence, and immune signaling, but with divergent patterns based on HA size and dose. Notably, supraphysiologic high-molecular-weight HA (HMW-HA) elicited a focused suppression of proliferative and inflammatory pathways consistent with a protective, quiescent phenotype, whereas low-molecular-weight HA (LMW-HA) induced broader, stress-associated changes. Unexpectedly, medium-molecular-weight HA (MMW-HA) exerted the strongest influence at physiologic concentrations, driving metabolic and cytoskeletal signaling. To our knowledge, this is the first study to apply global proteomics to dissect HA size- and dose-dependent effects on the lung endothelium, addressing a longstanding gap in the field.

HMW-HA has long been recognized as a homeostatic component of the extracellular matrix, supporting vascular integrity and dampening inflammation (Johnson et al., 2018; Garantziotis et al., 2018; Singleton, 2013). Consistent with this literature, our proteomic data show that HMW-HA at supraphysiologic dose suppresses cell cycle and inflammatory signaling, reinforcing the concept of HMW-HA as a “tissue integrity signal.” Prior studies in animal models of lung injury demonstrated that exogenous HMW-HA attenuates neutrophilic inflammation and protects barrier function (Johnson et al., 2018; Garantziotis et al., 2018), in part through CD44-mediated signaling and antagonism of Toll-like receptor 4 (TLR4) pathways (Taylor et al., 2004). Our finding that HMW-HA drives a senescent or quiescent endothelial phenotype extends these observations, suggesting that one mechanism by which HMW-HA stabilizes the endothelium is by actively limiting proliferative drive and immune activation. This interpretation is further supported by recent omics work showing that CD44–HA interactions at endothelial adhesion sites can concentrate HA and regulate leukocyte transmigration (van Steen et al., 2023) Moreover, plasma proteomics in septic shock patients revealed broad dysregulation of HA-associated proteins, particularly those involved in extracellular matrix stability (van der Heijden et al., 2025), underscoring that loss of intact HMW-HA networks is a feature of systemic inflammation. Together, these findings situate our results within a broader body of evidence that intact, supraphysiologic HMW-HA reinforces vascular stability by suppressing inflammatory and proliferative pathways.

In contrast, LMW-HA has consistently been described as a danger-associated molecular pattern (DAMP) that promotes inflammation (Jiang et al., 2011; Vistejnova et al., 2014). Fragments of HA generated during tissue injury can activate Toll-like receptors (TLR2 and TLR4) and CD44, leading to NF-κB activation and induction of cytokines such as IL-6, TNFα, and IL-1β (Taylor et al., 2004; Vistejnova et al., 2014).

Our proteomic data align with this paradigm: LMW-HA altered the largest number of endothelial pathways overall, but with diffuse, stress-associated signatures rather than focused suppression. This broad remodeling is consistent with prior work demonstrating that LMW-HA fragments drive leukocyte recruitment and endothelial activation in sterile lung injury models (Muto et al., 2019). Recent clinical studies further underscore this role: in patients with ARDS and COVID-19, elevated circulating HA, particularly fragmented forms, correlated with impaired oxygenation and higher mortality ( Li et al., 2024). Experimental work has further demonstrated that SARS-CoV-2 infection itself induces hyaluronan production in vitro and in vivo (Hellman et al., 2024), providing a mechanistic basis for the excessive HA accumulation observed in COVID-19 lungs.

Mechanistically, medium- and low-weight HA fragments are highly hygroscopic and can exacerbate alveolar edema by retaining water in the injured lung (Barnes et al., 2023). Thus, the diffuse, stress-like endothelial proteome we observed under LMW-HA treatment mirrors the pro-inflammatory, barrier-disruptive functions attributed to HA fragments in vivo.

In addition to the divergent effects of HMW- and LMW-HA, our data revealed a novel and unexpected role for medium-molecular-weight HA (MMW-HA). At physiologic concentrations (200 ng/mL), MMW-HA produced more robust proteomic changes than either HMW- or LMW-HA, prominently activating insulin secretion and Rho GTPase signaling pathways. This finding contrasts with the prevailing paradigm in which HA biology is often framed as a binary opposition between protective HMW-HA and pro-inflammatory LMW-HA, with little attention to intermediate forms.

Although understudied, MMW-HA has been reported to exert distinct bioactivity. In macrophages, HA fragments in the 250–1000 kDa range stimulate pro-inflammatory cytokine production (TNFα, IL-1β, IL-6, CCL2), similar to LMW-HA (Jiang et al., 2011; Cowman et al., 2015). In regenerative contexts, however, MMW-HA can enhance cell viability and tissue remodeling. A recent in vitro study found that both LMW- and MMW-HA significantly increased the metabolic activity and osteogenic differentiation of human periodontal ligament cells, whereas HMW-HA was less effective (Frasheri et al., 2023) Likewise, clinical work in tendinopathy has shown that ultrasound-guided injections of medium-weight HA reduce pain and improve tendon architecture, with decreased thickness and neovascularization at follow-up (Fogli et al., 2017). These observations suggest that MMW-HA may occupy a functional niche, balancing the reparative and stimulatory properties of LMW-HA with some of the stabilizing features of HMW-HA.

Our proteomic data extend this concept by showing that MMW-HA, specifically at physiologic levels, is uniquely potent in modulating endothelial pathways related to metabolism and cytoskeletal organization. The pronounced effect on insulin signaling is particularly noteworthy, as endothelial insulin resistance contributes to vascular dysfunction in inflammatory states. The Rho GTPase family, in turn, is central to cytoskeletal dynamics and endothelial barrier regulation. Thus, our results suggest that intermediate-sized HA fragments, often generated during tissue turnover, may play underappreciated roles in maintaining endothelial homeostasis under basal conditions. Further work is needed to define the receptor interactions and signaling mechanisms by which MMW-HA exerts these effects, as well as to test whether these findings translate in vivo.

Beyond size-specific effects, our network-level analyses revealed a conserved HA-response program across all molecular weights and concentrations. Pathways related to cell cycle control, senescence, and immune modulation were consistently altered, suggesting that HA, regardless of size, engages a core set of endothelial processes.

However, the magnitude and direction of these changes depended strongly on dose and molecular weight. At supraphysiologic concentrations, HMW-HA produced robust suppression of proliferative and inflammatory pathways, while LMW-HA triggered broader, stress-associated responses. At physiologic levels, responses were more muted overall, but MMW-HA uniquely drove strong activation of metabolic and cytoskeletal signaling. Together, these findings highlight that both molecular weight and concentration are critical determinants of HA bioactivity in the endothelium.

These insights may help explain clinical observations linking HA abundance and fragmentation with outcomes in acute lung injury. Circulating HA levels are markedly elevated in patients with ARDS and COVID-19, and higher concentrations correlate with worse survival (Li et al., 2024) Autopsy and imaging studies have described HA-rich, viscous alveolar exudates in severe COVID-19, consistent with the hygroscopic nature of fragmented HA and its ability to exacerbate edema (Barnes et al., 2023). Our data suggest that the balance of HA sizes and concentrations may dictate whether the endothelium adopts a protective, quiescent state (driven by HMW-HA) or a stress-activated, barrier-disruptive state (driven by LMW- and MMW-HA).

These findings also have therapeutic implications. HA is already widely used in medicine as an injectable viscosupplement, wound healing adjunct, and cosmetic filler (Cowman et al., 2015; Costa et al., 2023). Recent reports highlight that HA formulations of different sizes may have distinct clinical effects: medium-weight HA injections have improved pain and tendon structure in tendinopathy (Fogli et al., 2017), and oral HA supplementation has shown benefits for skin hydration and joint comfort (Gao et al., 2023). In parallel, experimental strategies to enzymatically degrade excess HA with hyaluronidase and DNase have shown promise in reducing the thickness of airway secretions in COVID-19 ARDS (Porter et al., 2024). Our results provide a mechanistic framework for these clinical observations, underscoring that the therapeutic benefit or harm of HA likely depends on both molecular weight and dose. Specifically, supplementing HMW-HA may stabilize the endothelial barrier, while blocking signaling from LMW- and MMW-HA fragments could mitigate inflammatory injury.

Our study has several limitations. First, we used an immortalized endothelial cell line (HULEC-5a), which may not fully recapitulate the complexity of primary pulmonary endothelium or the in vivo microenvironment. Second, we focused on a single time point (24 hours), providing a static view of HA-induced changes; dynamic time-course studies will be needed to capture early versus late signaling events. Third, while global proteomics identified numerous pathways and networks, functional validation was not performed for specific pathways such as insulin signaling or Rho GTPase activity. Finally, the concentrations studied were designed to reflect physiologic versus supraphysiologic ranges, but intermediate doses may also reveal important biological transitions that were not captured here.

Future studies should address these limitations by testing HA responses in primary endothelial cells and animal models of lung injury. Time-resolved omics approaches could define the temporal sequence of HA signaling, while targeted experiments (e.g., barrier integrity assays, receptor knockdowns) will clarify the mechanisms driving MMW-HA’s unique activity. In parallel, integrating proteomics with transcriptomics and metabolomics may provide a more comprehensive systems-level understanding of HA’s impact on the pulmonary endothelium. Translationally, studies in patient cohorts with **ARDS or sepsis could examine whether the HA-responsive pathways identified here** align with circulating HA size distributions and clinical outcomes.

In conclusion, this study demonstrates that HA is not a passive structural molecule but an active regulator of endothelial signaling, with effects strongly dependent on both molecular weight and concentration. HMW-HA at high doses reinforced a protective, quiescent phenotype, whereas LMW-HA induced broad stress-like changes, and MMW-HA unexpectedly emerged as the most potent regulator at physiologic levels. Together, these findings define a conserved but context-specific HA-response program in the lung endothelium. By linking size- and dose-dependent HA biology to global proteomic signatures, our work provides new mechanistic insight into how HA fragmentation and accumulation contribute to lung injury, and highlights opportunities for tailoring HA-based therapies in disease contexts where endothelial dysfunction is central.

## Author Contributions

J.A.M., K.K., and S.Y. conceived and designed the research. K.K performed all sample preparation. J.A.M. operated the mass spectrometer. J.A.M., K.K, and S.Y. carried out all aspects of data analysis. J.A.M. and S.Y interpreted results. S.Y. drafted the manuscript. J.A.M, K.K., and S.Y. edited the manuscript. All authors discussed each stage of the manuscript and have approved the final version of the manuscript.

## Materials & Correspondence

Correspondence and material requests can be addressed to Dr. James A. Mobley.

